# Comparative study of enriched dopaminergic neurons from siblings with Gaucher disease discordant for parkinsonism

**DOI:** 10.1101/2024.02.25.581985

**Authors:** Ellen Hertz, Gani Perez, Ying Hao, Krystyna Rytel, Charis Ma, Martha Kirby, Stacie Anderson, Stephen Wincovitch, Kate Andersh, Tim Ahfeldt, Joel Blanchard, Yue Andy Qi, Grisel Lopez, Nahid Tayebi, Ellen Sidransky, Yu Chen

**Affiliations:** Medical Genetics Branch, National Human Genome Research Institute, National Institutes of Health, Bethesda, MD; Center for Alzheimer’s and Related Dementias (CARD), National Institute on Aging and National Institute of Neurological Disorders and Stroke, National Institutes of Health, Bethesda, MD, USA; National Human Genome Research Institute, National Institutes of Health, Bethesda, MD; Nash Family Department of Neuroscience at Mount Sinai, New York, NY, USA; Aligning Science Across Parkinson’s (ASAP) Collaborative Research Network, Chevy Chase, MD, 20815

**Keywords:** iPSC, dopaminergic neuron differentiation, *GBA1*, Parkinson disease, neomycin resistance, Gaucher disease, neurodegeneration

## Abstract

Inducible pluripotent stem cells (iPSCs) derived from patient samples have significantly enhanced our ability to model neurological diseases. Comparative studies of dopaminergic (DA) neurons differentiated from iPSCs derived from siblings with Gaucher disease discordant for parkinsonism provides a valuable avenue to explore genetic modifiers contributing to *GBA1*-associated parkinsonism in disease-relevant cells. However, such studies are often complicated by the inherent heterogeneity in differentiation efficiency among iPSC lines derived from different individuals. To address this technical challenge, we devised a selection strategy to enrich dopaminergic (DA) neurons expressing tyrosine hydroxylase (TH). A neomycin resistance gene *(neo)* was inserted at the C-terminus of the *TH* gene following a T2A self-cleavage peptide, placing its expression under the control of the *TH* promoter. This allows for TH+ DA neuron enrichment through geneticin selection. This method enabled us to generate comparable, high-purity DA neuron cultures from iPSC lines derived from three sisters that we followed for over a decade: one sibling is a healthy individual, and the other two have Gaucher disease (GD) with *GBA1* genotype N370S/c.203delC+R257X (p.N409S/c.203delC+p.R296X). Notably, the younger sister with GD later developed Parkinson disease (PD). A comprehensive analysis of these high-purity DA neurons revealed that although GD DA neurons exhibited decreased levels of glucocerebrosidase (GCase), there was no substantial difference in GCase protein levels or lipid substrate accumulation between DA neurons from the GD and GD/PD sisters, suggesting that the PD discordance is related to of other genetic modifiers.

## INTRODUCTION

Biologically relevant *in vitro* models of dopaminergic (DA) neurons are of significant scientific interest due to the pivotal role of these cells in common diseases like Parkinson disease (PD). Since the 1950’s, the progressive degeneration of DA neurons in the substantia nigra has been identified as a defining feature and the cause of the cardinal motor symptoms in PD [1, 2]. However, the cellular mechanisms responsible for this neuronal loss remain largely elusive. Previous experimental models only partially recapitulate this multifactorial disorder, whereas the role of genetic vulnerability is now being increasingly appreciated [3]. The most common known genetic risk factors for PD to date are variants in *GBA1,* the gene encoding glucocerebrosidase (GCase) [4, 5]. Approximately 5-10% of patients with PD carry *GBA1* variants [6–8]. While *GBA1*-associated PD is clinically indistinguishable from sporadic disease, it exhibits an earlier age of onset and a faster disease progression in a subgroup of patients [7, 9]. Biallelic *GBA1* mutations result in the rare lysosomal storage disease Gaucher disease (GD). Despite a significant reduction of GCase activity in patients with GD, most do not develop parkinsonism, suggesting that secondary factors contribute to disease development [10, 11].

Comparative studies of DA neurons differentiated from induced pluripotent stem cells (iPSCs) derived from individuals discordant for PD offer an opportunity to study underlying pathogenic mechanisms in the affected cell type, while preserving the permissive or resistant genetic backgrounds [12]. However, several challenges exist in such studies. Firstly, there is often limited available clinical information regarding the donor of the iPSCs lines used. Most studies reported only gender, age of onset if applicable, and age at sample collection, but lacked description of disease-relevant symptoms, clinical course, and/or long-term outcome, limiting stratification of clinically similar cell lines and interpretation of *in vitro* results. Secondly, despite progress in differentiation methods, inherent heterogeneity in DA neuron differentiation efficiency among iPSC lines and culture batches complicates the interpretation and reproducibility of results [13, 14]. This variability may be related to the donor‘s overall genetic background rather than the genotypes of research interest. Efforts have been focused on refining differentiation protocols with timely addition of signaling molecules mimicking *in vivo* embryonic development [14, 15], but critical events for dopaminergic patterning, such as the level of *Wnt* activation [16–18], seem to be an inherent cell line characteristic [19]. Lastly, the final fraction of DA neurons is often limited in the culture, mixed with proliferating neural progenitor cells as well as other types of cells [20]. Consequently, disease-related changes in DA neurons may sometimes be obscured by culture heterogeneity.

Various methods have been employed to overcome this heterogeneity. Optimization of differentiation conditions for each cell line has been suggested, but this is time-consuming and introduces potential biases [21]. Expression of lineage-determining transcription factors has been successful in generating glutamatergic-like neurons [22], but similar efficiency for inducing DA neurons has yet to be achieved, even with combinations of transcription factors together with exogenous patterning cues [23, 24]. The DNA cross-linker mitomycin-C (MMC) has been used to remove proliferating progenitor cells in differentiating DA neuron cultures, which can sometimes outnumber the postmitotic DA neurons in the culture [25, 26]. Purification of DA neurons expressing tyrosine hydroxylase (TH), the rate-limiting enzyme in catecholamine synthesis, has been achieved through fluorescence activated cell sorting (FACS) with immunolabelled neurons [27] and those genetically engineered to express fluorescent proteins concomitant with TH [28, 29]. The latter also enables readily identification of TH-expressing cells in live cell applications. In addition, a neomycin (*neo*) resistance gene driven by a *MAP2*-promotor has previously been inserted to the safe-harbor AAVS1 site in iPSCs and used for selection of DA neurons [30], despite MAP2 being a pan-neuronal marker. Recently, single-cell RNA-sequencing analysis of iPSC-derived DA neuron cultures from 95 patient-derived iPSC lines revealed substantial line-to-line variation [20], with an average of 80% of differentiated cells being MAP2 positive, and only 20% also expressing TH. This landmark study emphasizes the inherent differentiation heterogeneity among iPSC lines and limited yield of DA neurons in some, calling for methods to standardize DA neuron culture for comparative studies.

To investigate genetic modifiers influencing PD penetrance in GD patients, we compared DA neurons differentiated from iPSCs derived from three siblings. Two of three sisters were diagnosed with type 1 GD, sharing *GBA1* genotype N370S/c.203del+R257X (p.N409S/c.203del +p.R296X), and one of them later developed PD. Their unaffected sister carried no *GBA1* mutation and showed no signs of PD. To increase the yield of DA neurons and reduce the heterogeneity among differentiated cultures, we inserted a neomycin resistance gene (*neo*) into the *TH* locus using CRISPR/Cas9-based genetic engineering, placing its expression under the control of the endogenous *TH* promotor. The *neo* insertion had no impact on iPSC differentiation into TH+ DA neurons, and the expression of neomycin phosphotransferase II (Neo II), encoded by *neo*, was concurrent with TH. Successful DA neuron enrichment with geneticin selection was confirmed using immunocytochemistry, western blots, and flow cytometry. Cellular proteomics of the DA neuron cultures also demonstrated reduced variability only after geneticin selection.

Finally, a comprehensive analysis of the high-purity DA neurons revealed comparable levels of GCase and lipid substrates in GD and GD/PD DA neurons, suggesting potential roles for genetic modifiers in the PD discordance.

## RESULTS

### Clinical presentation of the siblings with Gaucher disease discordant for parkinsonism

Two sisters (HT707 and HT708) carrying the *GBA1* genotype N370S/c.203del+R257X (p.N409S/c.203del+p.R296X) were longitudinally monitored for non-neuronopathic GD (GD1) under a natural history study at the National Institutes of Health [31]. HT707 was diagnosed with GD at age 12, exhibiting skeletal involvement, anemia, thrombocytopenia, and organomegaly, leading to a partial splenectomy. Although enzyme replacement therapy (ERT) improved hematologic parameters and organ size, skeletal complications persisted, and pulmonary hypertension developed later. Assessment by a movement disorder specialist at ages 60 and 62 revealed no evidence of parkinsonism. Unfortunately, she died at age 63 due to metastatic sarcoma, with autopsy findings indicating no Lewy body pathology and normal basal ganglia. Her younger sister, HT708, presented with hepatosplenomegaly at age 7 and was subsequently diagnosed with GD. She exhibited thrombocytopenia and fatigue, both of which showed improvement upon ERT. At age 55, she developed PD, initially presenting with a left-hand tremor. Examination at age 56 revealed bradykinesia, rigidity, and a unilateral left pill-rolling tremor, all of which responded well to L-DOPA treatment. After six years of treatment, she experienced motor fluctuations, leading to the initiation of L-DOPA infusion. Eight years after her PD onset, her cognition appeared to be normal, as evidenced by a Montral Cognitive Assessment (MoCa) score of 26 points. Their middle sister, HT711, carries no *GBA1* mutation and has shown no signs of PD during examinations at ages 58 and 63. A summary of their clinical findings is provided in Fig. 1a. Skin fibroblasts were obtained from HT707 at age 60, HT708 at 56 (after PD diagnosis), and HT711 at 58. Reprogramed iPSCs showed expected morphology (Fig. 1b) with concordant expression of the pluripotency markers OCT3/4, Tra1-60 and Nanog (Fig. 1c, d).

**Figure 1.**
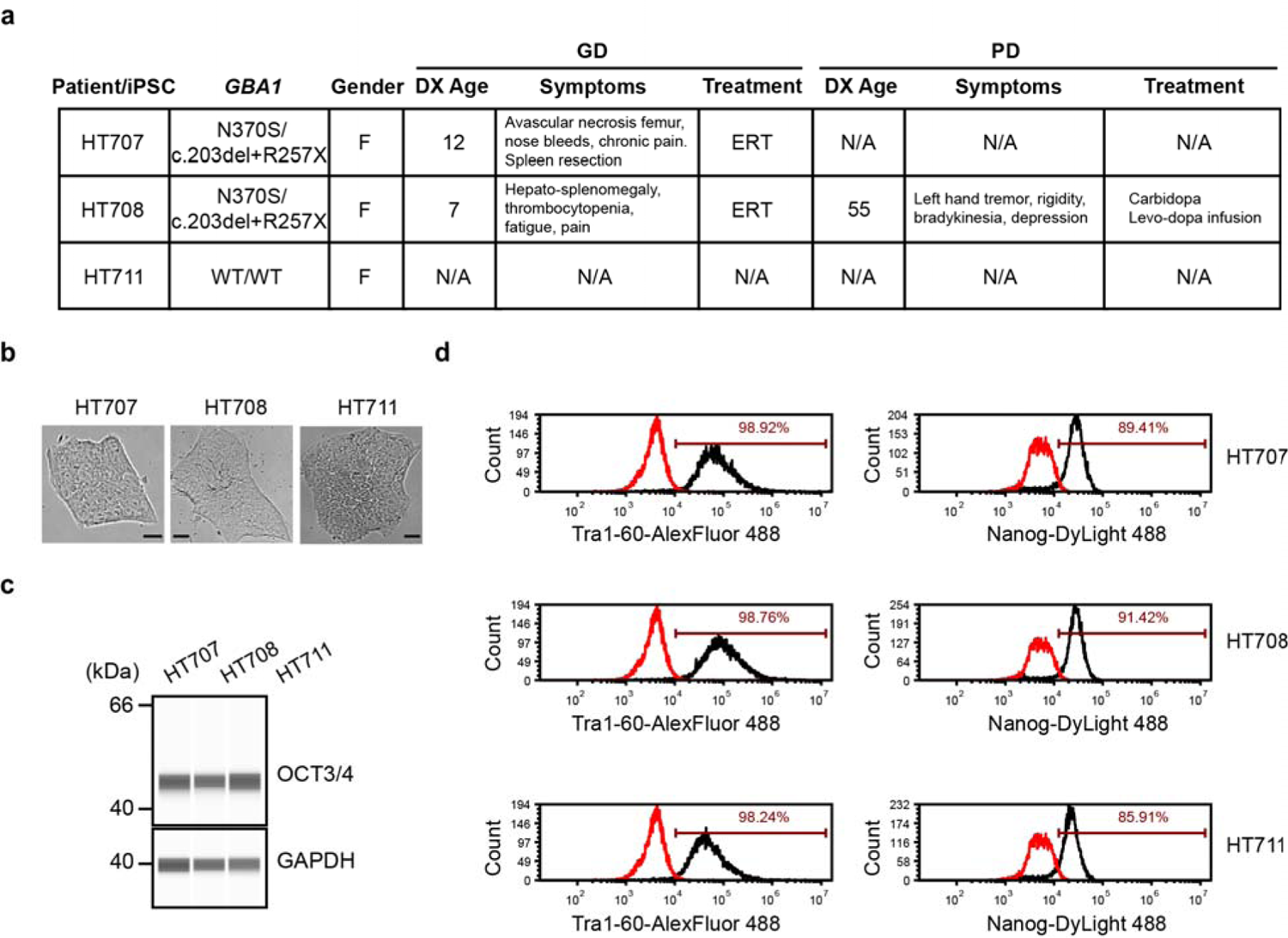
Characterization of patient-derived iPSC lines. **a** Summary of clinical features of iPSC donors. **b** Brightfield images of iPSCs (Scale bars, 10 µm) **c** The expression of pluripotency marker OCT3/4 in all iPSC lines demonstrated by Western blotting. **d** Quantification of the expression of pluripotency markers Tra1-60 and Nanog in all iPSC lines with flow cytometry.

### Genetic engineering of iPSC lines to enable DA neuron enrichment through geneticin selection

To address inherent variability in DA neuron differentiation among iPSC lines reprogrammed from different donors, we developed a method to enrich *TH*-expressing (TH+) DA neurons within iPSC-derived DA neuronal cultures. Employing CRISPR-Cas9 and homology-directed repair (HDR), we inserted a *neo* gene into the *TH* locus after exon 14, following a P2A self-cleavage peptide. Anticipating the expression of *neo,* driven by the endogenous *TH* promoter, to be restricted to TH+ DA neurons, we aimed to facilitate their enrichment with geneticin selection (Fig. 2a). Because the *TH* gene is transcriptionally silent in iPSCs, our HDR donor vector included an EF1α promoter-driven selection cassette. This cassette included a puromycin resistance gene and a red fluorescent protein, mRuby, separated by a T2A self-cleavage peptide. This design aimed to aid the identification of correctly edited iPSCs with puromycin selection and FACS sorting of mRuby+ iPSCs. Furthermore, the selection cassette was flanked by loxP sites to assist its removal after the edited iPSCs were identified. After the co-transfection of the HDR donor vector and the CRISPR plasmids encoding Cas9 protein and transcribing sgRNAs, iPSCs with HDR donor vector integration were enriched with puromycin selection. Enriched iPSCs were then transfected with a plasmid encoding GFP-tagged CRE-recombinase to remove the selection cassette. Individual iPSCs expressing CRE-GFP were sorted into 96-well plates via FACS to derive single cell clones. Edited iPSC clones were further analyzed using PCR to confirm the intended insertion. The edited iPSC clonal lines were referred to as HT707 *TH-neo*, HT708 *TH-neo*, and HT711 *TH-neo*, respectively. Their morphology remained unchanged and they exhibited normal karyotypes (Fig. 2b, c).

**Figure 2.**
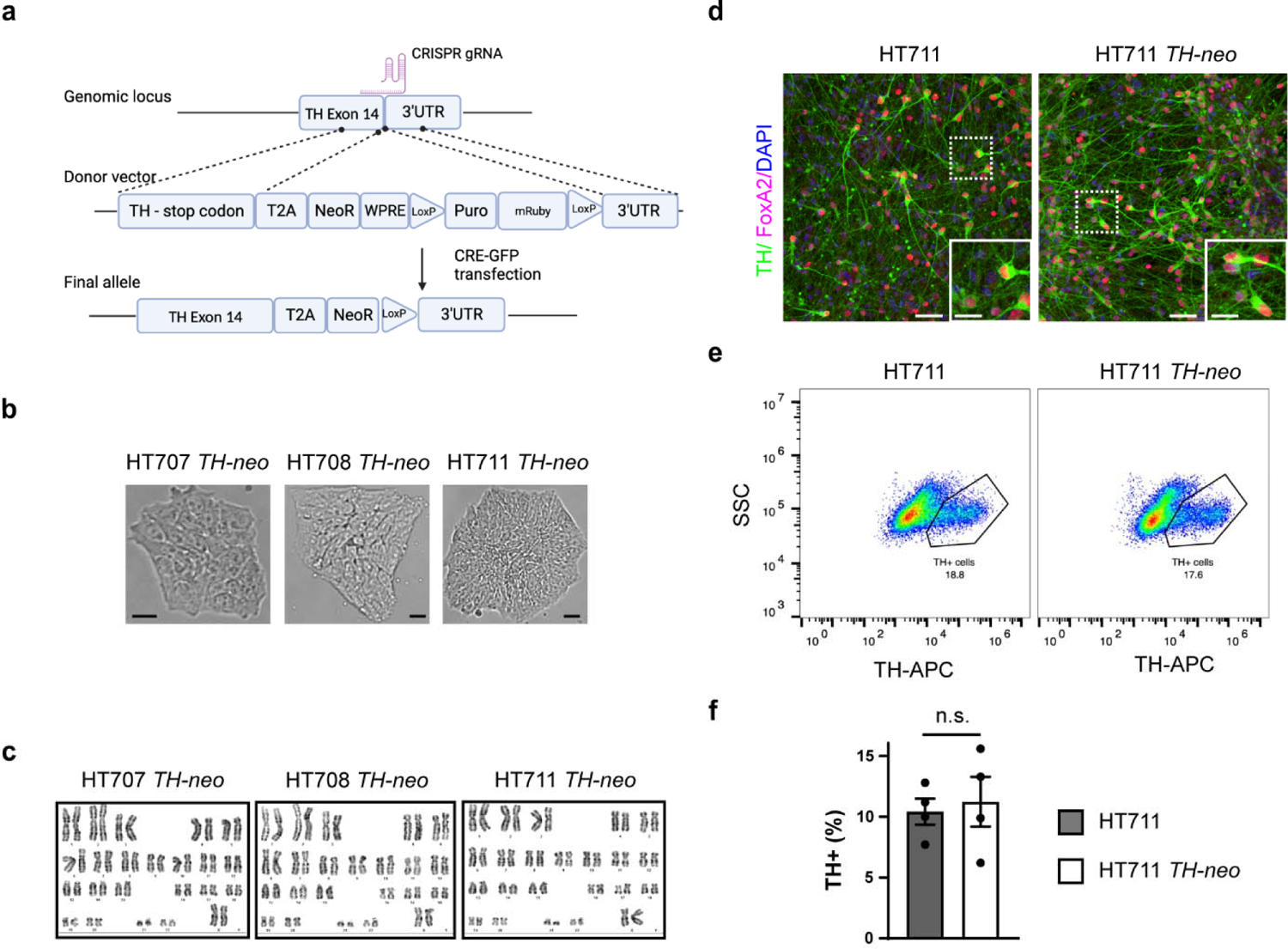
Generation and characterization of *TH-neo* edited iPSC lines. **a** CRISPR-mediated knock-in of *neo* in the TH locus. **b** Brightfield images of *TH-neo* iPSC lines (Scale bar, 10 µm). **c** Normal karyotypes of *TH-neo* iPSC lines. **d** Immunocytochemistry (ICC) of TH+ cells in HT711 and HT711 *TH-neo* at day 37 of differentiation (Scale bar, 50 µm; insert scale bar, 25 µm). **e,f** Quantification of TH+ cells using flow cytometry in HT711 and HT711 *TH-neo* at day 37 of differentiation (*t*-test, *n*=4 independent differentiations).

**Figure 3.**
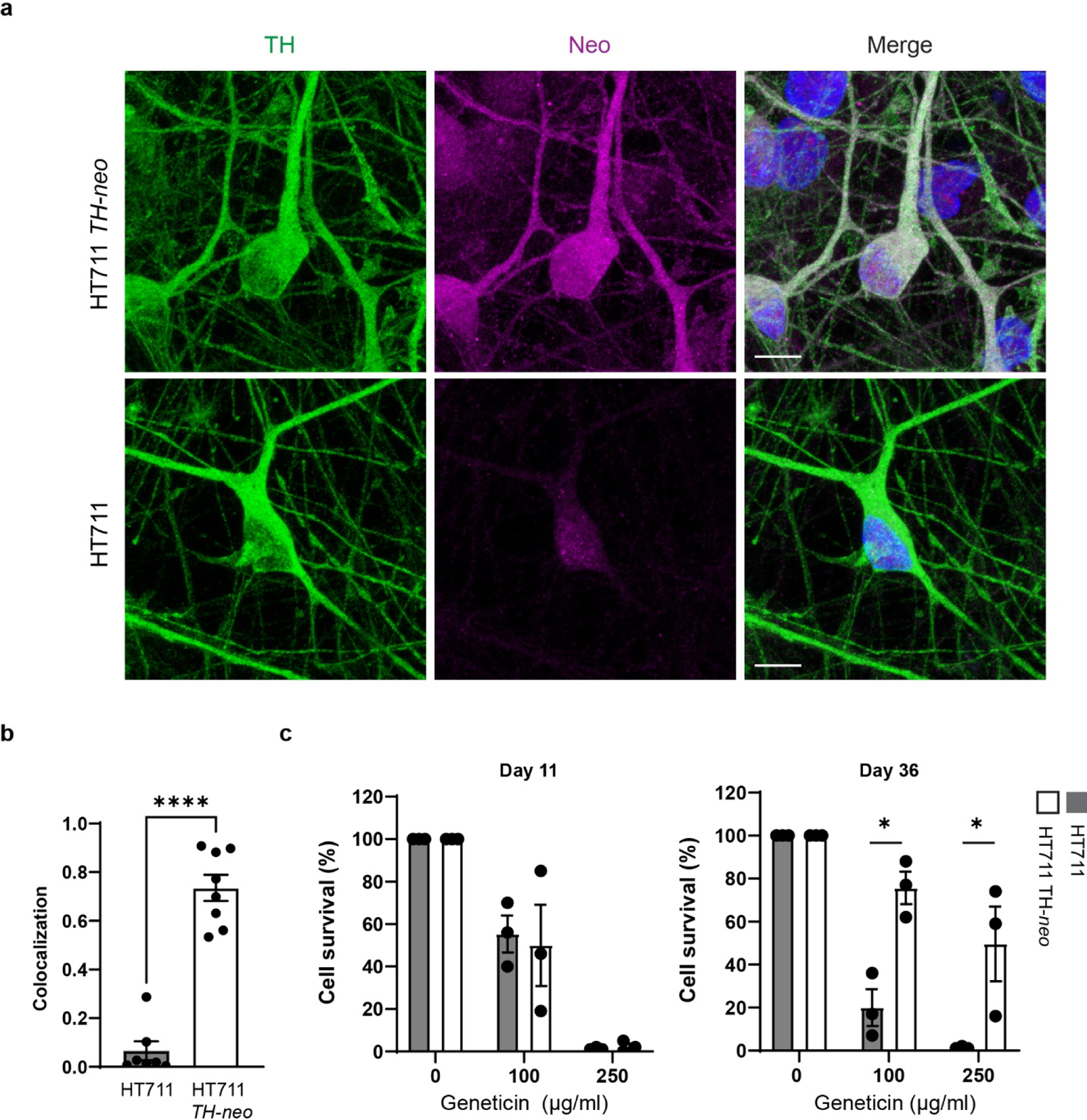
*neo* expression is limited to TH+ neurons. **a** The expression of *neo* is limited in TH+ neurons as demonstrated with ICC. **b** Quantification of colocalization. No correlation is detected in HT711 (*t*-test, ***, *p*<0.001, *n*=3 independent differentiations). **c** Geneticin eliminates both HT711 and HT711 *TH-neo* neural progenitor cells (*n*=3 independent differentiations). **d** Dose dependent survival of HT711 *TH-neo* DA neurons following geneticin treatment (ANOVA, \*\**p*=0.01, \*\*\**p*<0.001, *n*=3 independent differentiations).

### Integration of the *neo* gene does not interfere with the differentiation of iPSCs into DA neurons

To evaluate the potential influence of *neo* gene integration on iPSC differentiation into DA neurons, HT711 and HT711 *TH-neo* were differentiated into DA neurons following a protocol developed by Kim et. al [16]. The differentiation efficiency, measured by the percentage of TH+ DA neurons at day 37 of differentiation, were comparable between HT711 and HT711 *TH-neo*, as evaluated by immunofluorescence and flow cytometry. This indicates that *neo* gene integration did not perturb the differentiation capacity (Fig.2d-f).

### Concurrent expression of tyrosine hydroxylase and neo in TH+ DA neurons

For efficient enrichment of DA neurons through geneticin selection, it is crucial for the *neo* gene to be expressed exclusively in TH+ DA neurons. Immunostaining on day 37 of differentiation revealed precise expression of neomycin phosphotransferase II (Neo II), the protein encoded by *neo*, exclusively in TH+ DA neurons differentiated from HT711 *TH-neo*. This indicates the expression of Neo II was under the control of the TH promotor (Fig. 2a, b). In contrast, TH+ DA neurons from HT711 showed minimal background staining, confirming the specificity of the Neo II antibody. To validate geneticin selection stress on proliferating neuronal precursors and neurons lacking TH and Neo II, we evaluated cell viability at both the neuronal precursor and neuronal stages using CellTiter-Glo luminescent cell viability assay. At the neuronal precursor stage (day 16), prior to the onset of TH expression, geneticin treatment similarly reduced the viability of HT711 and HT711 *TH-neo* neuronal precursors (Fig. 2c). At a concentration of 250 μg/ml, geneticin effectively eliminated neuronal precursors, demonstrating its efficacy in removing proliferating precursors from DA neuron cultures. When assessed at day 37 of neuronal differentiation, after 10 days of treatment with 250 μg/ml geneticin, ∼50% of the cells in HT711 *TH-neo* DA neuron cultures survived, whereas nearly all cells in HT711 DA neuron cultures perished, indicating successful elimination of neurons not expressing Neo II through geneticin selection (Fig. 2d).

### Geneticin treatment increases the fraction of TH+ DA neurons

To illustrate the enrichment of TH+ DA neurons after geneticin selection, we quantified the percentage of TH+ DA neurons using both immunofluorescence and flow cytometry analyses. Immunofluorescence studies revealed that treatment with 250 ug/ml of geneticin increased the fraction of TH+ cells from 10% to 35% (Figs. 4a, b), while flow cytometry demonstrated an increase from 10% to 45% (Fig. 4e). Furthermore, western blot analysis exhibited a dose-dependent increase in the expression of TH following geneticin selection, with an approximately 2.5-fold increase observed at a concentration of 250 μg/ml (Fig. 4c).

**Figure 4.**
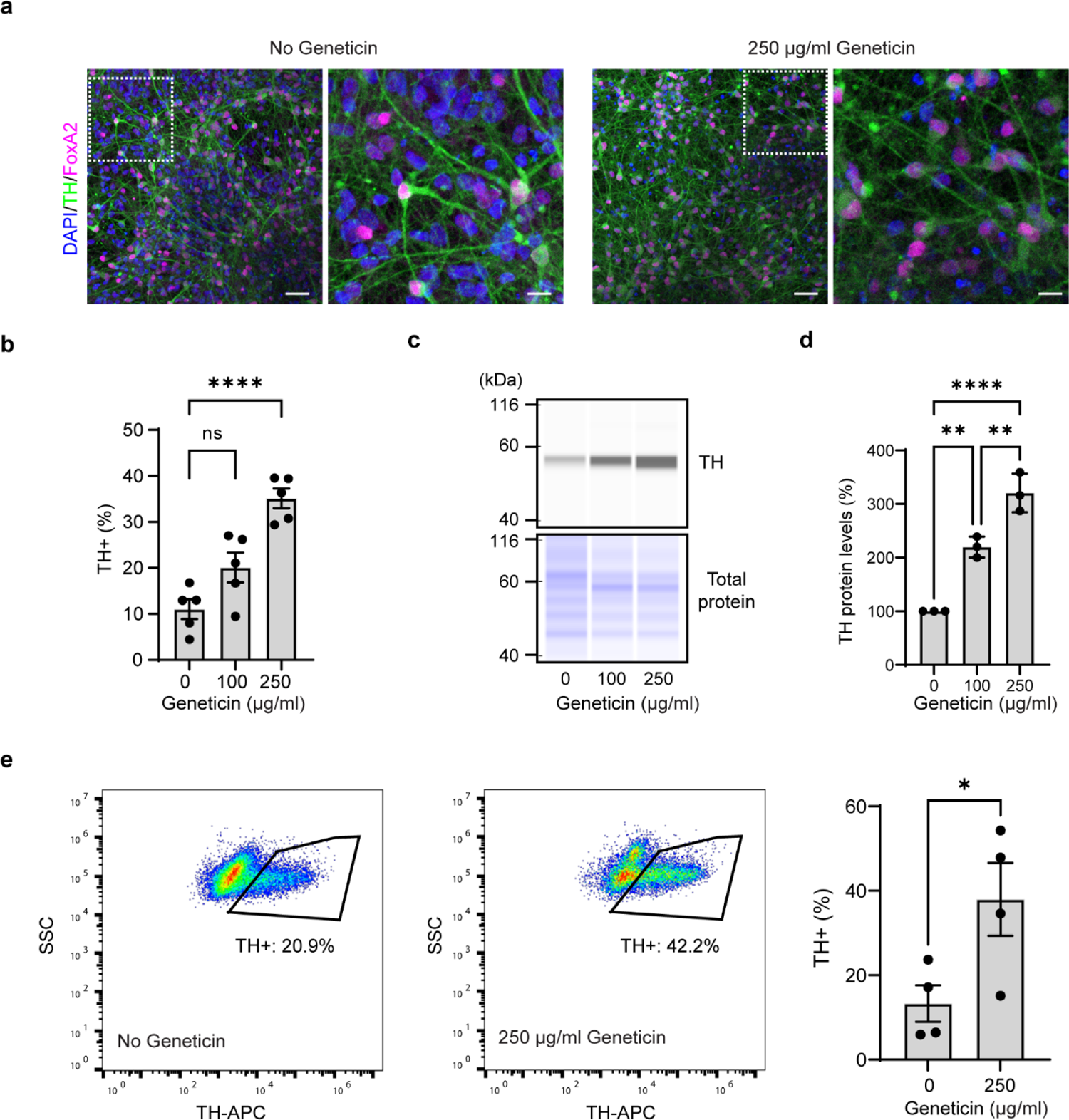
Enrichment of TH+ neurons with geneticin treatment. **a** Confocal image of TH+ DA neurons with and without geneticin treatment. Note the reduction of the nuclei without accompanying TH staining. **b** Quantification of TH+ DA neurons in ICC (1-way ANOVA, \*\*\**p*<0.001, *n*=5 fields of view from 2 independent differentiations). **c, d** Western blot analysis of TH expression. Data is normalized to cells without geneticin treatment (1-way ANOVA, **p <0.01, *n*=3 independent differentiations). **e** Flow analysis of TH+ cells. (t-test, \**p*<0.05, *n*=4 independent differentiations).

### Standardization of DA neurons across patient-derived iPSC lines was confirmed by cellular proteomics

The primary goal of the novel tool was to attain comparable yields of TH+ DA neurons from the three patient-derived iPSC lines, facilitating meaningful genotype and genetic background comparisons, avoiding confusion from variable differentiation efficiency. To assess the cellular composition in DA neuron culture pre- and post-geneticin selection, we employed a robust proteomics pipeline involving automated sample preparation and data-independent acquisition (DIA) coupled with a high-field asymmetric waveform ion mobility spectrometer (FAIMS) interface, and an optimized library-free DIA database search strategy [32]. We treated the cultures with geneticin or vehicle for 10 days and continued to mature them to day 75 for cellular proteomics analysis. The pipeline detected over 8800 proteins, indicating a comprehensive coverage of the cellular proteome.

All three cell lines exhibited an increased abundance of TH after the geneticin treatment (711 *TH-neo* log2FC 2.14, adj. p.<1×10^-3^, 707 *TH-neo* log2FC 0.70, adj. p.<2.5×10^-5^, 708 *TH-neo* log2FC 1.22, adj.p <3×10^-6^) (Fig. 5a, Fig. S1). Additionally, enhanced expression of other DA neuronal markers was evident, including dopa decarboxylase (DDC), which coverts L-DOPA to dopamine (711-*TH-neo* log2FC 2.14, adj. p.<8×10^-5^, 707-*TH-neo* log2FC 0.68, adj. p.<1×10^-4^, 708-*TH-neo* log2FC 1.75, adj.p 5×10^-7^), and ALDH1A1, a specific marker for PD-vulnerable A9 DA neurons (708-*TH-neo* log2FC 1.60, adj. p.<0.002, 711-*TH-neo* log2FC 2.32, adj. p.=0.013). In contrast, the pan-neuronal markers MAP2 and TUBB3 were not significantly enriched. This is consistent with the previous report that a large proportion of iPSC-derived neurons were MAP2 positive but lacked TH expression [20].

**Figure 5.**
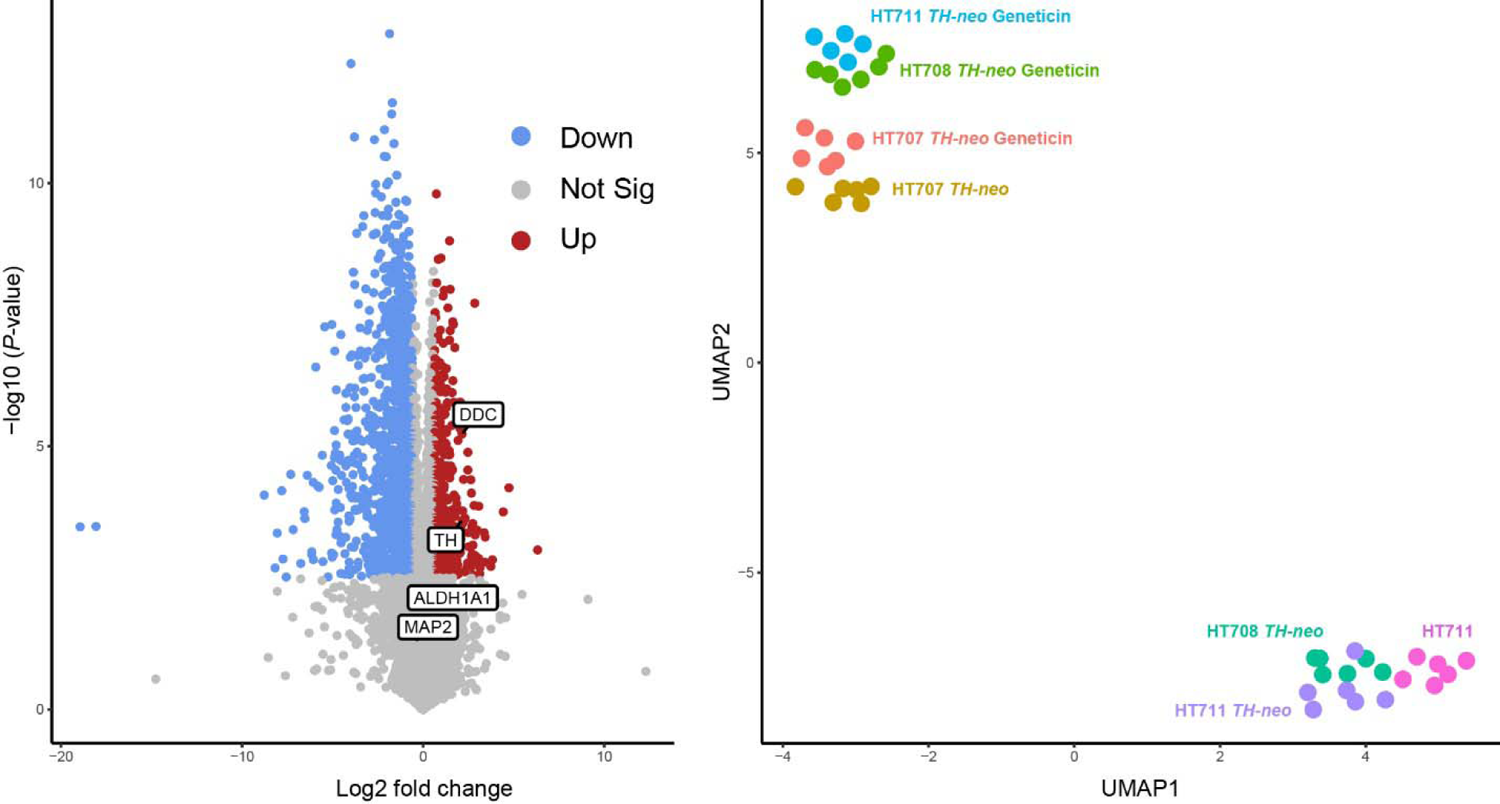
Cellular proteomics comparing DA neurons. **A** Volcano plot showing differentially expressed proteins in HT711 *TH-neo* DA neurons with and without geneticin selection. DA neuron markers enriched by geneticin treatment include TH, DDC and ALDH1A1 (*n*=*1* independent differentiation). **b** UMAP clustering of HT707, HT708, and HT711 DA neurons shows improved comparability after geneticin selection.

Dimensionality reduction by Uniform Manifold Approximation and Projection (UMAP) was then used to visualize overall similarities between DA neuron cultures derived from the cell lines. HT711 and HT711 *TH-neo* clustered together, confirming that individual characteristics were maintained in the edited cell lines. HT707 *TH-neo* with and without geneticin selection shared similar proteome profiles, indicating geneticin selection did not have major impact on the cellular proteome. Prior to geneticin selection, HT708 *TH-neo* and HT711 *TH-neo* clustered together with the parental line HT711. Geneticin selection led to substantial increases in the levels of DA markers in HT708 *TH-neo* and HT711 *TH-neo*, and their cellular proteomes moved closer to HT707 *TH-neo*. Collectively, geneticin selection reduced the overall heterogeneity among the three DA neuron cultures and set the stage for further phenotyping (Fig. 5b).

### Phenotyping high purity DA neuron cultures

The western blot analysis of enriched DA neurons from the three siblings revealed comparable TH levels in all cultures, confirming the successful harmonization of the cultures. In contrast, TH levels exhibited greater variability in the cultures differentiated from the unedited parental lines, highlighting differentiation disparities among iPSC lines and underscoring the necessity for standardization (Fig. 6a). GCase was found to be downregulated in HT707 and HT708 compared to HT711, with similar levels between HT707 and HT708. This suggests *GBA1* genotypes has a more substantial impact on GCase levels than the PD status (Fig. 6b). Lipidomic analysis of GlcCer and GlcSph, through supercritical fluid high-performance liquid chromatography/mass spectrometry (SFC/LC-MS/MS) revealed no substantial accumulation of GlcCer or GlcSph in HT707 or HT708 DA neurons (Fig. 6c).

**Fig 6.**
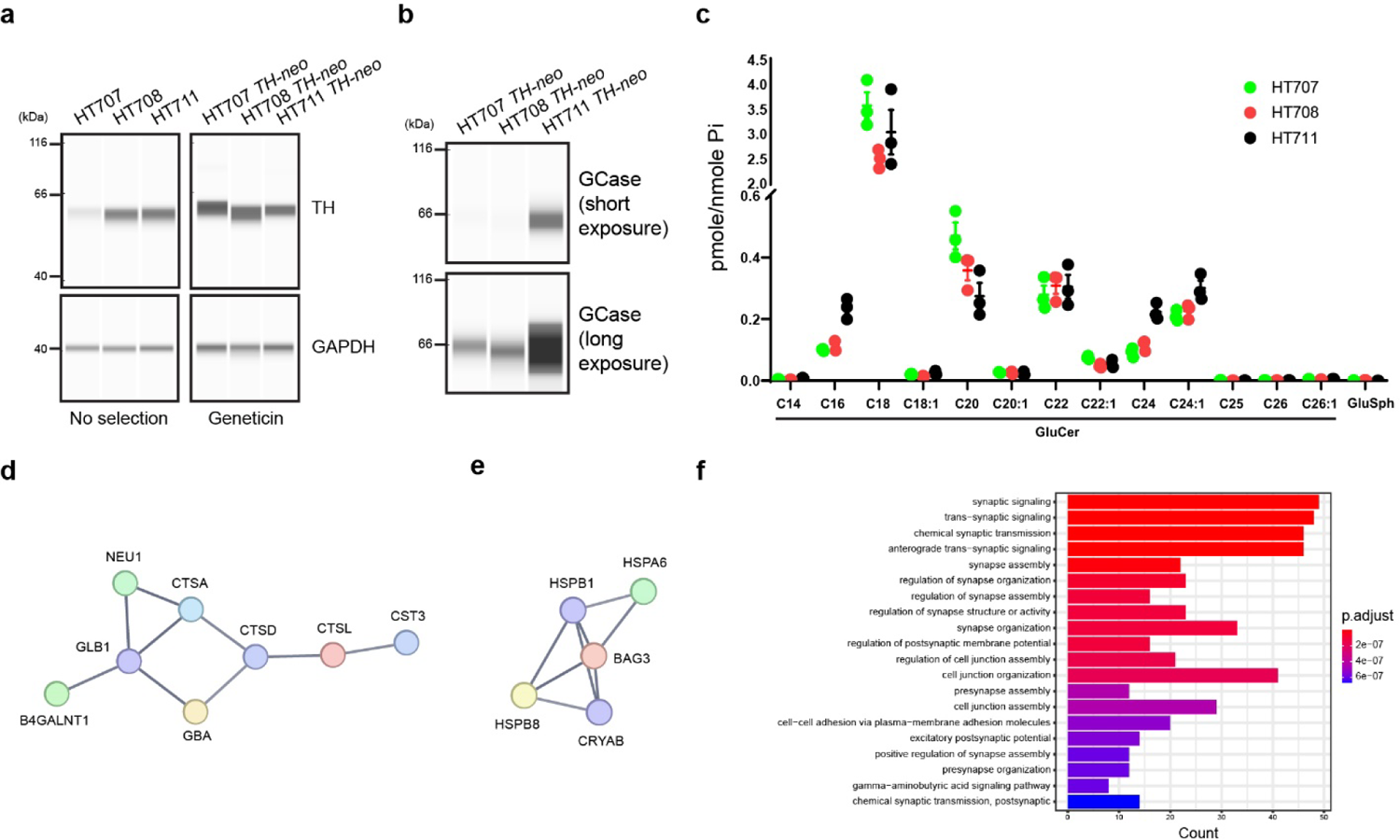
Phenotyping high-purity DA neurons. **a,b** Western blotting analyzing enriched DA neurons from HT707 *TH-neo*, HT708 *TH-neo*, and HT711 *TH-neo* as well as those from the parental lines. **c** Lipidomics analysis of enriched DA neurons. **d** A network of lysosomal hydrolases that is down-regulated in DA neurons from both HT707 *TH-neo* and HT708 *TH-neo*. **e** A network of molecular chaperones that is upregulated in HT708 *TH-neo* DA neurons compared to HT707 *TH-neo*. **f** Top 20 enriched GO term pathways comparing HT708 *TH-neo* and HT707 *TH-neo* DA neurons.

To characterize neuronal changes linked to non-neuronopathic GD, we delved deeper into the cellular proteome dataset and identified shared changes in HT707 *TH-neo* and HT708 *TH-neo* compared to HT711 *TH-neo*. This included 80 upregulated and 118 downregulated proteins (cut-off: fold change 1.5, adj. P-value <0.01.). Protein-Protein interaction networks functional enrichment analysis using the STRING database (https://string-db.org/) revealed a network of lysosomal proteins among the shared downregulated proteins, encompassing *GBA1*, *CTSA*, *CTSL* and *GLB1* (Fig. 6d), indicating broad lysosomal changes as a consequence of GD. To delineate neuronal changes specifically linked to PD, we investigated the differences between HT707 *TH-neo* and HT708 *TH-neo.* A network of molecular chaperones, including HSPA6 of the heat shock 70 kDa protein family and its co-chaperone BAG3, were upregulated in HT708 *TH-neo*, potentially suggesting disruptions in protein homeostasis due to the PD genetic background (Fig. 6e). GO term enrichment analysis of proteomics data also indicated major differences in synapse assembly between HT707 *TH-neo* and HT708 *TH-neo*, identifying “synaptic signaling” and “synapse assembly” among the top downregulated biological (Fig. 6f).

## Discussion

Thorough assessments of rare or unique clinical cases offer valuable insights into underlying molecular mechanisms [33, 34]. iPSCs hold great promise as models for better recapitulating individual-specific molecular signatures in disease-relevant cell types. Here, we present three iPSC lines from a family with a genetic predisposition for PD, yet exhibiting disparate clinical outcomes, along with associated longitudinal clinical data. To maximize the benefits of patient-derived models, a detailed understanding of the clinical trajectory is crucial, allowing for correlations to *in vitro* findings, stratification of iPSC lines based on clinical phenotype, and potentially informing treatment responses [35]. In this family, comprehensive clinical information, including a post-mortem neuropathology examination, confirmed that HT707 had no indication of parkinsonism, despite harboring biallelic *GBA1* mutations resulting in low expression of GCase in *in vitro* DA neurons, unlike her affected sister. Linking this clinical information to the three genetically related iPSC-lines provides a unique model that can potentially shed light on factors contributing to or protecting against the development of *GBA1*-related PD.

To facilitate accurate downstream comparisons, we addressed the substantial differentiation heterogeneity inherent in iPSC lines, even among lines derived from genetically related individuals. Previous advancements, including optimized differentiation protocols for specific iPSC lines from healthy controls, deep phenotyping of selected cell lines with the desired stem cell characteristics [36], and data atlases generated from numerous iPSCs in large-scale collaborative projects [20], may not be applicable when studying rare diseases or atypical clinical cases, as in our case. Consequently, we aimed to develop a tool capable of enriching the desired cell type, thereby reducing expected heterogeneity between patient-derived samples. After placing the expression of *neo* under the control of the TH promoter, we first verified the exclusive expression of *neo* in TH+ DA neurons. The effectiveness of geneticin selection was then demonstrated by comparing cell survival between the parental line and the corresponding *TH-neo* line. Virtually no parental cells survived the 250 μg/ml geneticin treatment, confirming the efficacy of this dose for eliminating neuronal precursor cells and neurons not expressing Neo II. Subsequently, we showed the enrichment of TH+ neurons using multiple methods. Immunostaining confirmed an increased faction of TH+ neurons post geneticin selection, caused by a decrease in DAPI-stained nuclei lacking TH. Flow cytometry analysis also revealed comparable enhancement. Compared to immunostaining, flow cytometry analysis required similar hands-on time, but analyzed a much larger number of neurons, thereby further strengthening our quantification. To comprehensively characterize the cellular composition post-selection, we generated cellular proteomics datasets and visualized their clustering using UMAP [37]. It was revealed that DA neuron cultures from the three patient iPSC lines clustered together only after enrichment. This emphasizes the critical importance of comparable differentiation efficiency for uncovering disease-relevant phenotypes through refined comparisons between individuals. This new method, complimentary to the previous approaches such as coupling a fluorescent reporter to TH expression, offers the advantage of cell enrichment without the need for dissociation into single cells, and can also be applied at any point of maturation after the onset of TH expression.

Finally, we characterized the enriched DA neurons from the three sisters. Despite the expected finding that the DA neurons from HT707 and HT708 DA exhibited lower GCase levels than HT711, no substantial accumulation of GCase lipid substrates was found in either line. This aligns with their diagnosis of non-neuronopathic GD, where no GD-related neuronal pathology and little accumulation of lipid substrates are observed in in post-mortem brain tissues[38, 39]. Comparable GCase and lipid substrate levels in HT707 and HT708 implicate the role of other genetic modifiers in the PD discordance.

In summary, we show that coupling geneticin resistance to endogenous TH expression can be used to harmonize dopaminergic cultures from different individuals, serving as an approach particularly suitable for the evaluation of rare clinical samples.

## Methods

### Obtaining and genotyping fibroblasts and reprogramming into iPSCs

The three participants were evaluated under the IRB-approved clinical protocol 86HG0096 at the National Institutes of Health, and each provided informed consent prior to participation. Sanger sequencing of the glucocerebrosidase gene was performed as previously described[40]. Fibroblasts were obtained by punch biopsy and cultured in Dulbecco’s Modified Eagle’s Medium (DMEM, 10566016; Thermo Scientific) supplemented with 15% fetal bovine serum (FBS; G22133, R&D) supplemented with 1× penicillin-streptomycin (Cat.# 15140-122; Thermo Scientific) at 37 °C. The CytoTune™-iPS 2.0 Reprogramming System (Cat.# A16517; Thermo Fisher), based on non-integrative Sendai virus, was used to generate iPSCs at the National Heart, Lung and Blood Institute’s iPSC Core according to standard procedures.

### Genomic engineering of iPSCs

iPSCs were maintained in Essential 8 Medium (Cat.#A1517001 Thermo Fisher) on Vitronectin-coated plates (Cat.#A14700; Thermo Fisher) without addition of antibiotics. Media was exchanged daily, and cells were kept at 37 °C and 5% CO^2^. Cells were passaged by dissociation in 0.5 mM EDTA every 3-4 days and resuspended in Essential 8 Medium supplemented with Y-27632 dihydrochloride (10 μM; Cat.#1254; R&D Systems). For CRISPR-Cas9 editing, iPSCs were dissociated to single cells with StemPro-Accutase (Cat.#A1110501; Gibco) and seeded in Essential 8 Medium with Y-27632 at a density of 200,000 cells/cm^2^ two hours prior to transfection with Lipofectamine Stem Transfection Reagent (Cat.#STEM00001; Invitrogen). A donor plasmid was generated, similar to that previously described by Ahfeldt et al. [41], with a 5’ TH homology arm, a 2A self-cleaving peptide sequence followed by *neo* - cassette and a WPRE sequence. In addition, a floxed selection cassette including a puromycin resistance cassette and mRuby sequence, and a 3’ TH homology arm (Fig. 1A), was co-transfected with vectors expressing gRNA sequences targeting the second to last codon of the human TH-locus (excluding the stop codon) and Cas9. Cells were replated the next day in StemFlex (Cat.#A3349401; Thermo Scientific), supplemented with Y-27632 and 24 hours later selection with 0.25 μg/ml puromycin (Cat.# A11138-03, Thermo Scientific) was initiated. Puromycin-resistant clones were allowed to replicate until near confluence and then pooled. CRE-mediated excision of the selection cassette was performed as described above, and single clones were collected by FACS on a Sony SH800 cell sorter the following day. Gates were determined by non-transfected control iPSCs. Single clones were sorted into a laminin-521 coated plate with Stemflex supplemented with 10 μM Y-27632, 5 μM Emricasan (Cat.# 7310, R&D), 1× penicillin-streptomycin, 0.7 μM ISRIB (Cat.# 5284, R&D) and 1× polyamines (Cat.# P8483, Sigma) and incubated for 3 days. Then, half of the media was changed with Stemflex supplemented with penicillin-streptomycin, ISRIB and polyamines. After this, the media was changed every three days with Stemflex alone, until the surviving clones reached at least 50% confluency and could be expanded.

Clones were screened for the correct knock-in by PCR, amplifying both cleavage sites. DNA was extracted from flash frozen pellets using 30 μl of QuickExtract (Lucigen # QE09050) and samples were heated to 60 °C for 15 min, 65 °C for 15 min and 99 °C for 10 min. AmpliTaq Gold 360 Master Mix (Thermo Fisher, #4398881) were mixed 1:2 with H_2_O, 0.25 μM of forward and reverse primers and 1 μL of cell extract. The primer sequences and PCR cycle conditions can be found in Supplement 1.

### Karyotyping

iPSC clones, which were confirmed for knock-in of the construct, were dissociated and plated as described above at 1×10^6^ cells/well in 6 well plates in Stemflex with 10 μM Y-27632. The following afternoon, Colcemid (1:100, Cat.# 10295892001, Roche) was added, and cells were karyotyped using standard procedures at the National Human Genome Research Institute Cytogenetics Core Facility.

### Directed differentiation into midbrain dopamine neurons (mDA)

The differentiation protocol for dopaminergic neurons was adopted from Kim et. al. with minor adaptations [16]. In short, iPSCs were dissociated into single cells with Accutase and plated on Geltrex-coated plates (Cat.#A1413302, Gibco) at 400,000 cells/cm^2^ in Neurobasal Medium (Cat.#21103-049) with 1× GlutaMAX (Cat.# 35050079, Gibco), 1× N2 (Cat.#17502048, Thermo Scientific), 1× B-27 without Vitamin A (Cat.#12587010, Thermo Scientific) supplemented with 500 ng/ml human Sonic Hedgehog/Shh (C24II) N-Terminus (Cat.# 1845-SH-500, R&D), 250 nM LDN 193189 dihydrochloride (Cat.# 6053, R&D), 2 μM A 83-01 (Cat.# 2939, R&D), 0.7μM CHIR99021(Cat.# 4423, R&D), SAG 21k (Cat.# 5282, R&D), 10μM Y-27632 and 5 μM Emricasan. On day 4, CHIR99021 was increased to 3 μM. On days 8 and 9, cells were cultured in Neurobasal/GlutaMAX/1X N2/B-27 with only 3 μM of CHIR99021. On day 10, media was changed to Neurobasal/GlutaMAX/B-27 with 20 ng/ml brain-derived neurotrophic factor (BDNF, Cat.# 450-02, Peprotech), 20 ng/ml glial cell line-derived neurotrophic factor (GDNF, Cat.# 450-10, Peprotech), 200 μM ascorbic acid (Cat#4034, Sigma-Aldrich), 1 ng/ml transforming growth factor type β3 (TGFβ3, Cat.# 243-B3, R&D), 200 μM dibutyryl cAMP (X, Sigma-Aldrich) and 3 μM CHIR99021. On day 11 cells were sub-plated at 800,000 cells/cm^2^ with Accutase in the same media with added Y-27632 and Emricasan. On day 13, CHIR99021 was substituted with 10 μM DAPT (Cat.# 2634, R&D). Cells were dissociated for the last time at day 25 and seeded at 200,000 cells/cm^2^. On day 27, selection with Geneticin/GENETICIN (Cat.# 10131-035, Gibco) was initiated and kept for 10 days until experiments were conducted on day 37, unless otherwise specified. For experiments conducted at day 75 of differentiation, media was changed once per week with the addition of 1 μM 5-Fluoro-2′-deoxyuridine (Cat.#F0503, Sigma-Aldrich) and 1 μM Uridine (Cat.#U3003, Sigma-Aldrich) to prevent division of non-differentiated cells.

### Cell viability assay – Cell-Glo

Cells were seeded in white opaque 96-wells with clear bottom after regular dissociation on day 11 and day 25, respectively and treated with Geneticin. On day 19 or 37, CellTiter-Glo (Cat# G7570, Promega) was resuspended with buffer and added 1:1 vol/vol directly to the media and placed 2 min on an orbital shaker. After 10 min incubation at RT in the dark, luminescence was detected on Flexstation 3 (Molecular devises).

### Western blots – automated system

Dopaminergic neurons and iPSC pellets were resuspended in Triton-X (1% Triton X-100, 10% glycerol, 150 mM NaCl, 25 mM HEPES pH 7.4, 1 mM EDTA, 1.5 mM MgCl2, supplemented with one EDTA-free protease inhibitor tablet per 10 mL of buffer (Pierce, cat no A32965)) or RIPA (Cat.# 20-188, Millipore) buffer, lysed through thorough pipetting or 3 freeze-thaw cycles and centrifuged at 21,000 X g for 15 minutes at 4°C. The supernatant was transferred to new tubes. Protein concentration was determined by BCA (Cat.# 23225, Thermo Scientific) according to manufacturer’s instructions and absorbance values were detected on Flexstation 3 (Molecular devises).

Comprehensive instructions for Jess Automated Western Blot System (Cat#. 004-650 BioTechne) are found with the kit and were run according to instructions in chemiluminescent RePlex mode with EZ Standard Pack 1 (Cat # PS-ST01EZ-8, BioTechne). In brief, the DTT was prepared with 40 μl of deionized water and the biotinylated ladder with 20 μl of deionized water respectively. Fluorescent 5X Master Mix was prepared with 20 μl of Sample Buffer and 20 μl of the DTT solution. Samples were prepared by combining one part fluorescent master mix with four parts sample to a final concentration of 1 μg/μl. Samples and a biotinylated ladder were incubated for 5 min at 95 °C. Fluorescence Separation Capillary Cartridges (12-230 kDa, Cat. # SM-W001, BioTechne) were loaded according to instructions in the RePlex pack (Cat# RP-001, BioTechne) with 3 μL sample per well. Primary antibodies (details in supplementary table 2) were diluted in in antibody diluent 2 and normalization was based on total protein or reference gene. Plates were centrifuged for 5 min at 1000 X g before addition of wash buffer, luminol-peroxide mix and RePlex reagent. Visualization was performed with SW Compass.

### Flow cytometry

Neurons were washed in PBS (Cat.# 14190144, Thermo Fisher) and incubated with Membrite (Cat.# 30093, Biotium) prestain solution, diluted 1:1000 in HBSS, for 5 min at 37 °C. Prestain solution was changed to staining solution and incubated for another 5 min at 37 °C. Cells were dissociated with prewarmed papain (Cat.#LK003176, Worthington) with DNAse (Cat.#LS006333, Worthington), Y-27632 and Emricasan for 10 min at 37°C. The reaction was quenched in DMEM/Glutamax (Cat.#10566-016, Thermo Scientific) with 10% fetal bovine serum and collected by centrifugation at 400Xg for 5 min. The pellet was resuspended in 4% paraformaldehyde (Cat.# sc-281692, Santa Cruz Biotechnology) and incubated for 10 min at RT on a rolling wheel. Cells were resuspended in DMEM with 10% FBS and spun at 500Xg for 5 min after which the cells were incubated in permeabilization buffer (1% BSA, 0.1% saponin in PBS, 10% normal donkey serum (Cat.# AB_2337258, Jackson ImmunoResearch) for 30 min at RT and thereafter labeled with tyrosine hydroxylase antibody or isotype controls (1:400, Cat.# P40101-150, Pel-Freeze and Cat.# AB-105-C R&D respectively) overnight at 4 °C. Cells were washed twice with 2% BSA and 1.5 mM EDTA in PBS and then incubated with diluted Alexa647-conjugated secondary antibody (Cat.# 711-605-152, Jackson ImmunoResearch), for 1 h at RT. Finally, cells were washed as described above with NucBlue Fixed Cell ReadyProbes Reagent (DAPI) (Cat. #R37606, Thermo Scientific) in the second wash and resuspended in wash buffer for flow cytometry analysis on a Beckman Coulter Cytoflex-LX. Initial gating included Membrite-positive cells, thereafter using DAPI positivity in order to exclude duplicates. Finally, the cells were analyzed for the fluorescent intensity of TH using Flow-Jo.

### Immunocytochemistry (ICC)

Cells were seeded in Geltrex-coated 8-well imaging chambers (Cat# 80807-90, Ibidi) at differentiation day 25. After selection as described above, cells were fixed in 4% paraformaldehyde (15 min at RT) and washed 3×5 min in PBS on day 37. Thereafter, cells were blocked for 1h at RT and incubated overnight with primary antibodies diluted in blocking buffer (1% BSA, 0.1% saponin in PBS) at 4 °C. The primary antibodies used were tyrosine hydroxylase (1:400, Cat.# P60101-150, Pel-Freeze) and FoxA2 (Cat.#RA34008, Neuromics). The following day, cells were carefully washed 3×5 min with PBS and incubated with secondary antibodies (Cat.# 711-545-152, 711-605-152, Jackson ImmunoResearch) for 1 h at RT. Cells were washed as described above, with NucBlue™ Fixed Cell ReadyProbes™ Reagent (DAPI) (Cat #R37606) included in the second set, and imaged in PBS. Confocal images were acquired on a Zeiss LSM 880 microscope with a 20x or 63x objective. Quantification of the TH-positive fraction was done with a custom-made program in IPP11.3.

### Proteomics

Dopaminergic neurons from the three patient-derived cells lines were differentiated in parallel and treated with 250 μg/ml GENETICIN or vehicle at differentiation day 27-37. Thereafter, differentiation media including uridine and FdU was changed weekly. On day 74/75 of differentiation, cells were washed three times in pre-chilled PBS and a lysis buffer (50 mM Tris-HCI, 50 mM NaCl, 1% SDS, 1% Triton X-100, 1%NP-40, 1% Tween 20, 1% glycerol, 1% Sodium deoxycholate (wt/vol), 5 mM EDTA) supplemented with 5mM Dithiothreitol (DTT), Benzonase and protease inhibitor were added just before collection. Samples were heated to 65°C at 1200 rpm for 30 min prior to the addition of iodoacetamide IAA (10 mM Cat#. A39271, Thermo Scientific). Samples were stored at −80 °C until analysis. Six technical replicates were analyzed for each condition. The proteomics pipeline including automated data analysis was applied as previously described [32, 42]. Protein-protein interaction networks was analyzed with the STRING database (https://string-db.org/) with the high confidence setting. Volcano plots were generated in Galaxy (https://galaxyproject.org/).

### Lipidomic analysis

Dopaminergic neurons were differentiated as described above but treated with 100 μg/ml GENETICIN at differentiation day 27-37 and the matured until day 75. Lipidomic analysis was performed by the lipidomics core facility at the Medical University of South Carolina.

Glucosylceramide, glucosylsphingosine, and galactosylceramide were analyzed by supercritical fluid high-performance liquid chromatography/ mass spectrometry (SFC/LC-MS/MS). Lipids were normalized to inorganic phosphate levels and run in technical triplicates.

---

**Supplementary Figure 1.**
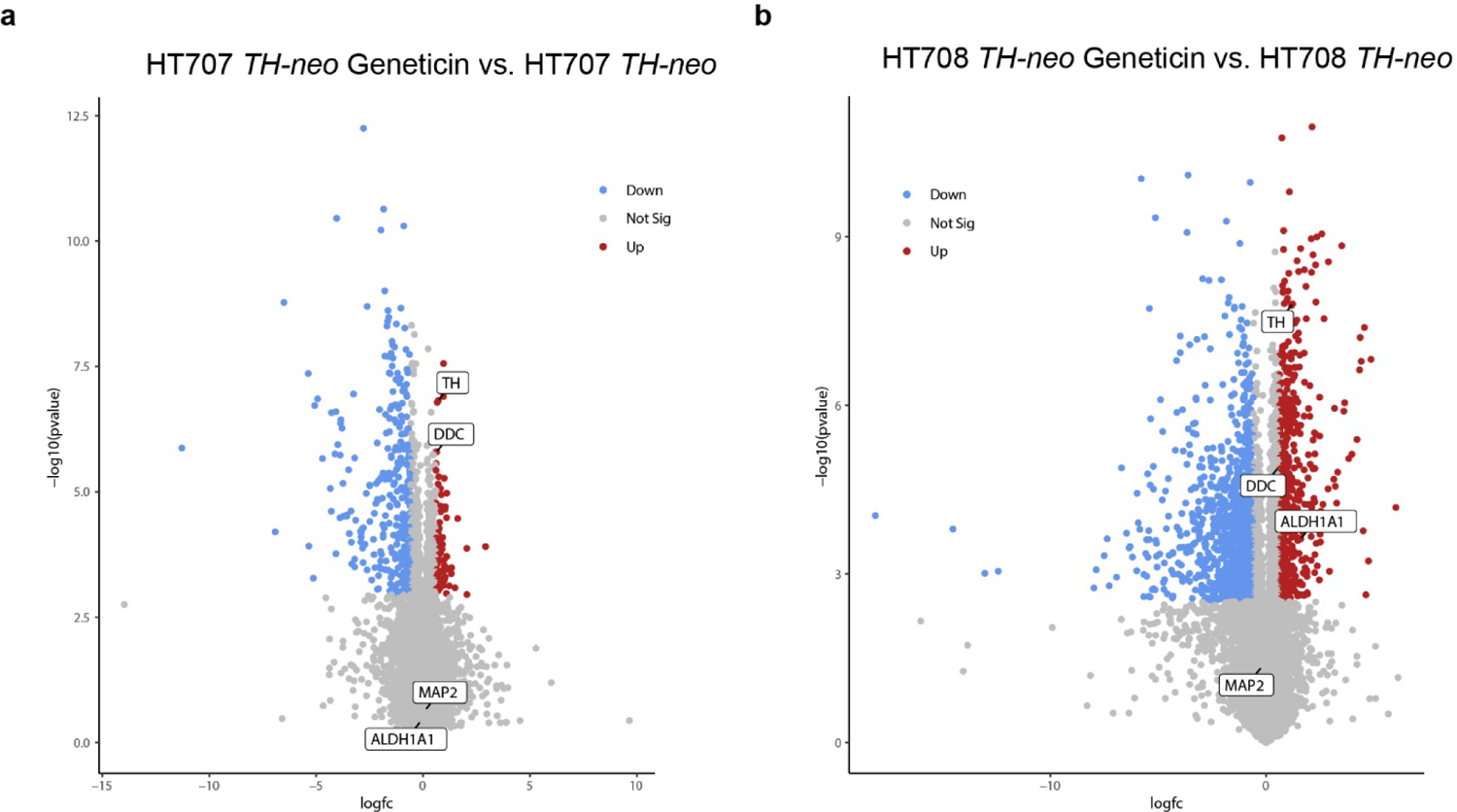
Volcano plots showing enrichment of dopaminergic markers for HT707 *TH-neo* (a) and HT708, *TH-neo* (b) after treatment with geneticin.

**Supplementary Table 1.**
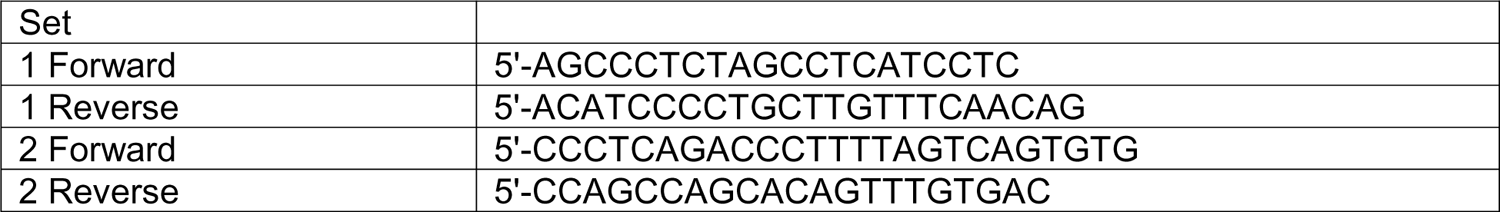
Primers used.

**Supplementary Table 2.** Antibodies used in this study.

| Antibody | Company | Catalogue number | Application |
| --- | --- | --- | --- |
| GAPDH | abcam | Ab9485 | WB (1:2000) |
| GCase | Abnova | H00002629-M01 | WB (1:50) |
| TH | Pel-Freeze | P60101-150 | IF (1:400), FC (1:400), WB (1:50) |
| FoxA2 | Neuromics | RA34008 | IF (1:500) |
| Neo II | Sigma | 06-747 | IF (1:500) |

## References

1. Carlsson, A., M. Lindqvist, and T. Magnusson, 3,4-Dihydroxyphenylalanine and 5-hydroxytryptophan as reserpine antagonists. Nature, 1957. 180(4596): p. 1200.

2. Hornykiewicz, O., Dopamine (3-hydroxytyramine) and brain function. Pharmacol Rev, 1966. 18(2): p. 925–64.

3. Nalls, M.A., et al., Identification of novel risk loci, causal insights, and heritable risk for Parkinson’s disease: a meta-analysis of genome-wide association studies. Lancet Neurol, 2019. 18(12): p. 1091–1102.

4. Sidransky, E., et al., Multicenter analysis of glucocerebrosidase mutations in Parkinson’s disease. N Engl J Med, 2009. 361(17): p. 1651–61.

5. Nalls, M.A., et al., A multicenter study of glucocerebrosidase mutations in dementia with Lewy bodies. JAMA Neurol, 2013. 70(6): p. 727–35.

6. Anheim, M., et al., Penetrance of Parkinson disease in glucocerebrosidase gene mutation carriers. Neurology, 2012. 78(6): p. 417–20.

7. Cilia, R., et al., Survival and dementia in GBA-associated Parkinson’s disease: The mutation matters. Ann Neurol, 2016. 80(5): p. 662–673.

8. Gan-Or, Z., et al., Genotype-phenotype correlations between GBA mutations and Parkinson disease risk and onset. Neurology, 2008. 70(24): p. 2277–83.

9. Lerche, S., et al., Dementia with lewy bodies: GBA1 mutations are associated with cerebrospinal fluid alpha-synuclein profile. Mov Disord, 2019. 34(7): p. 1069–1073.

10. Alcalay, R.N., et al., Comparison of Parkinson risk in Ashkenazi Jewish patients with Gaucher disease and GBA heterozygotes. JAMA Neurol, 2014. 71(6): p. 752–7.

11. Rosenbloom, B., et al., The incidence of Parkinsonism in patients with type 1 Gaucher disease: data from the ICGG Gaucher Registry. Blood Cells Mol Dis, 2011. 46(1): p. 95–102.

12. Shi, Y., et al., Induced pluripotent stem cell technology: a decade of progress. Nat Rev Drug Discov, 2017. 16(2): p. 115–130.

13. Yamanaka, S., Pluripotent Stem Cell-Based Cell Therapy-Promise and Challenges. Cell Stem Cell, 2020. 27(4): p. 523–531.

14. Kee, N., et al., Single-Cell Analysis Reveals a Close Relationship between Differentiating Dopamine and Subthalamic Nucleus Neuronal Lineages. Cell Stem Cell, 2017. 20(1): p. 29–40.

15. Nishimura, K., et al., Single-cell transcriptomics reveals correct developmental dynamics and high-quality midbrain cell types by improved hESC differentiation. Stem Cell Reports, 2023. 18(1): p. 337–353.

16. Kim, T.W., et al., Biphasic Activation of WNT Signaling Facilitates the Derivation of Midbrain Dopamine Neurons from hESCs for Translational Use. Cell Stem Cell, 2021. 28(2): p. 343–355 e5.

17. Kriks, S., et al., Dopamine neurons derived from human ES cells efficiently engraft in animal models of Parkinson’s disease. Nature, 2011. 480(7378): p. 547–51.

18. Oosterveen, T., et al., Pluripotent stem cell derived dopaminergic subpopulations model the selective neuron degeneration in Parkinson’s disease. Stem Cell Reports, 2021. 16(11): p. 2718–2735.

19. Blauwkamp, T.A., et al., Endogenous Wnt signalling in human embryonic stem cells generates an equilibrium of distinct lineage-specified progenitors. Nat Commun, 2012. 3: p. 1070.

20. Bressan, E., et al., The Foundational Data Initiative for Parkinson Disease: Enabling efficient translation from genetic maps to mechanism. Cell Genom, 2023. 3(3): p. 100261.

21. Gantner, C.W., et al., An Optimized Protocol for the Generation of Midbrain Dopamine Neurons under Defined Conditions. STAR Protoc, 2020. 1(2): p. 100065.

22. Zhang, Y., et al., Rapid single-step induction of functional neurons from human pluripotent stem cells. Neuron, 2013. 78(5): p. 785–98.

23. Ng, Y.H., et al., Efficient generation of dopaminergic induced neuronal cells with midbrain characteristics. Stem Cell Reports, 2021. 16(7): p. 1763–1776.

24. Sheta, R., et al., Combining NGN2 programming and dopaminergic patterning for a rapid and efficient generation of hiPSC-derived midbrain neurons. Sci Rep, 2022. 12(1): p. 17176.

25. Hiller, B.M., et al., Mitomycin-C treatment during differentiation of induced pluripotent stem cell-derived dopamine neurons reduces proliferation without compromising survival or function in vivo. Stem Cells Transl Med, 2021. 10(2): p. 278–290.

26. Zambon, F., et al., Cellular alpha-synuclein pathology is associated with bioenergetic dysfunction in Parkinson’s iPSC-derived dopamine neurons. Hum Mol Genet, 2019. 28(12): p. 2001–2013.

27. Sandor, C., et al., Transcriptomic profiling of purified patient-derived dopamine neurons identifies convergent perturbations and therapeutics for Parkinson’s disease. Hum Mol Genet, 2017. 26(3): p. 552–566.

28. Uberbacher, C., et al., Application of CRISPR/Cas9 editing and digital droplet PCR in human iPSCs to generate novel knock-in reporter lines to visualize dopaminergic neurons. Stem Cell Res, 2019. 41: p. 101656.

29. Calatayud, C., et al., CRISPR/Cas9-mediated generation of a tyrosine hydroxylase reporter iPSC line for live imaging and isolation of dopaminergic neurons. Sci Rep, 2019. 9(1): p. 6811.

30. Wakeman, D.R., et al., Cryopreservation Maintains Functionality of Human iPSC Dopamine Neurons and Rescues Parkinsonian Phenotypes In Vivo. Stem Cell Reports, 2017. 9(1): p. 149–161.

31. Lopez, G., et al., Clinical evaluation of sibling pairs with gaucher disease discordant for parkinsonism. Mov Disord, 2020. 35(2): p. 359–365.

32. Reilly, L., et al., A fully automated FAIMS-DIA mass spectrometry-based proteomic pipeline. Cell Rep Methods, 2023. 3(10): p. 100593.

33. Singleton, A.B., et al., alpha-Synuclein locus triplication causes Parkinson’s disease. Science, 2003. 302(5646): p. 841.

34. Lwin, A., et al., Glucocerebrosidase mutations in subjects with parkinsonism. Mol Genet Metab, 2004. 81(1): p. 70–3.

35. Rivetti di Val Cervo, P., et al., hiPSCs for predictive modelling of neurodegenerative diseases: dreaming the possible. Nat Rev Neurol, 2021. 17(6): p. 381–392.

36. Pantazis, C.B., et al., A reference human induced pluripotent stem cell line for large-scale collaborative studies. Cell Stem Cell, 2022. 29(12): p. 1685–1702 e22.

37. Winchester, L., et al., Comparing multiple clustering approaches to understand proteomic datasets for improved biomarker detection: Developing topics. Alzheimer’s & Dementia, 2020. 16: p. e047654.

38. Wong, K., et al., Neuropathology provides clues to the pathophysiology of Gaucher disease. Mol Genet Metab, 2004. 82(3): p. 192–207.

39. Tayebi, N., et al., Gaucher disease with parkinsonian manifestations: does glucocerebrosidase deficiency contribute to a vulnerability to parkinsonism? Mol Genet Metab, 2003. 79(2): p. 104–9.

40. Tayebi, N., et al., Genotypic heterogeneity and phenotypic variation among patients with type 2 Gaucher’s disease. Pediatr Res, 1998. 43(5): p. 571–8.

41. Ahfeldt, T., et al., Pathogenic Pathways in Early-Onset Autosomal Recessive Parkinson’s Disease Discovered Using Isogenic Human Dopaminergic Neurons. Stem Cell Reports, 2020. 14(1): p. 75–90.

42. Li, Z., et al., ProtPipe: A Multifunctional Data Analysis Pipeline for Proteomics and Peptidomics. bioRxiv, 2023.

